# imAgeScore, a Cell Painting-Based Predictor of Cellular Age for High-throughput Drug Screening Applications

**DOI:** 10.64898/2026.03.06.710056

**Authors:** Eleni Patili, Fabio M D’Orazio, Jack Brelstaff, Koby Baranes, Rodrigo L dos Santos, Mark RN Kotter, Joana M Tavares

## Abstract

Quantifying cellular age in vitro in a scalable and biologically meaningful manner is essential for the discovery of pharmacological interventions that modulate aging. We developed imAgeScore, a machine-learning model trained on high-content Cell Painting features to predict the phenotypic age of primary human dermal fibroblasts. imAgeScore correlates with chronological and DNA methylation–based age estimates and captures coordinated morphological changes across nuclear and cytoplasmic compartments. The model detected age acceleration during serial propagation and age reduction following partial reprogramming. Pharmacological interventions targeting distinct aging hallmarks induced predictable shifts in predicted age and enabled classification of damaging versus rejuvenating cellular states. Application of imAgeScore in an automated high-throughput screening pipeline identified candidate age-modulating compounds, revealed inter-individual variability in response magnitude, and detected additive rejuvenation effects in selected combinatorial treatments. Functional validation in a scratch wound assay confirmed enhanced cellular repair by leading candidates, supporting the biological relevance of morphology-derived age reduction. Together, these results demonstrate that image-based morphological profiling provides a scalable platform for quantifying cellular aging and screening for candidate rejuvenation interventions.

## Introduction

Aging is a major risk factor for many chronic diseases, and interventions that slow or reverse age-associated cellular decline hold substantial promise for improving health span. While genetic and environmental factors that modulate aging have been extensively studied ^1–4^, there remains a critical need for pharmacological compounds that can induce rejuvenation effects in human cells. Identifying such compounds requires experimental systems that can sensitively and quantitatively measure changes in cellular age, enabling the systematic discovery and evaluation of interventions with true rejuvenation potential.

Cellular aging is characterized by a diverse set of molecular and functional changes, often described in terms of distinct “hallmarks of aging”, which are categorized into primary (genomic instability, telomere attrition, epigenetic alterations, loss of proteostasis), antagonistic (deregulated nutrient sensing, mitochondrial dysfunction, cellular senescence) and integrative (stem cell exhaustion, chronic inflammation, altered intercellular communication) hallmarks^5,6^. Despite their utility, interpreting aging in vitro remains challenging when it relies on separately assessing multiple hallmarks^6–8^. Each hallmark typically requires a distinct assay, often involving different experimental protocols, readouts, and time scales^9^. This approach is labor-intensive, costly, and difficult to integrate into a single interpretable measure of cellular age or health. Moreover, many hallmark-specific assays are low-throughput and not readily scalable to automated screening workflows^6,7^. As a result, there is a need for models that capture cellular age in vitro in a way that is both biologically meaningful and operationally compatible with drug discovery.

Several molecular aging clocks have been developed to quantify cellular age in vitro, most notably epigenetic and transcriptomic clocks, but both present limitations in screening contexts. Epigenetic clocks based on DNA methylation patterns are among the most accurate predictors of chronological age^10,11^. However, these assays remain relatively expensive and low-throughput, and they primarily capture epigenetic alterations corresponding to a single hallmark of aging, making them poorly suited for large-scale discovery of compounds that modulate cellular aging. Transcriptomic aging clocks, by contrast, estimate cellular age through the measurement of thousands of transcripts in a more hallmark-agnostic manner and can predict chronological age and age-related risk with high accuracy^12–16^. Although transcriptomic data accurately reflects biological states, it does not allow direct conclusions about cell function. Despite their biological richness, transcriptomic approaches remain costly in practice, particularly at the scale required for compound screening.

An alternative approach is image-based phenotypic profiling, which uses high-content microscopy to measure multiple cellular parameters simultaneously at single-cell resolution^17,18^. Cell Painting (CP) is a standardized implementation of image-based profiling that facilitates multiplexed fluorescent labelling of key cellular structures, enabling the detection of a vast range of phenotypes across experimental conditions^19–21^, which often reflect cellular function. The Cell Painting assay was designed to be simple and inexpensive to implement in high-throughput screening facilities, relying exclusively on fluorescent dyes. Following staining and imaging, automated analysis pipelines such as Harmony and CellProfiler^22^ extract thousands of morphological features, making Cell Painting a scalable and unbiased assay for genetic and compound screening as well as mechanism-of-action studies^23–26^.

Here, we present a machine-learning model that predicts the phenotypic age of fibroblasts in vitro from high-content imaging data acquired using a Cell Painting assay. Rather than relying on predefined hallmark-specific readouts, the model integrates diverse image-derived features that collectively capture cellular state, yielding a hallmark-unbiased estimate of fibroblast age compatible with high-throughput drug discovery workflows. We applied this approach to human skin fibroblasts derived from 24 donors spanning a wide age range. Using extracted morphological features, we derived a phenotypic aging signature, termed imAgeScore, which correlated with both chronological donor age and experimentally validated epigenetic age. We further demonstrated that imAgeScore detects age-associated changes in fibroblasts that were serially propagated until senescence^27^, as well as age reversal following transient reprogramming^28^. Finally, we validated imAgeScore’s ability to capture aging- and rejuvenation-associated phenotypes induced by known modulatory compounds and applied it to screen pre-assembled libraries of drugs and supplements to identify candidate rejuvenation compounds and novel combinations.

## Methods

### Cell Culture

Primary normal human dermal fibroblasts (NHDFs) derived from 24 donors were obtained from multiple commercial suppliers. Donors spanned a broad age range (0–87 years) and included both sexes, multiple ethnicities and different biopsy sites to account for potential confounding variables. From this point onward, donors are denoted by their age at the time of biopsy (in years) and sex, where M denotes male and F denotes female (e.g. 44F indicates a 44 years old female donor) (Supplementary Table 1). Cells were cultured on 0.1% gelatin–coated dishes (StemCell Technologies, 7903) and maintained in DMEM/F-12 (Thermo Fisher Scientific, 21331020) supplemented with 10% fetal bovine serum (FBS; Sigma, F9665-500ML), non-essential amino acids (Thermo Fisher Scientific, 11140050), GlutaMAX (Thermo Fisher Scientific, 35050061), 2-mercaptoethanol (Thermo Fisher Scientific, 31350010), and 1% penicillin–streptomycin (Gibco, 15140122). Culture medium was replaced every two days.

Original vials were expanded to generate a cell bank (see Supplementary Methods). For subsequent experiments, banked vials were thawed at passage 2 and cultured for four days before seeding into 96-well PhenoPlates (Revvity, 6055302) at a density of 5,000 cells per well. Cells were cultured for an additional three days prior to Cell Painting or other staining procedures.

### Propagation model generation

For propagation experiments, NHDFs from donors 0M, 4M, 22F, 44F, and 62F were thawed and continuously cultured in the presence of FGF2. Cells were serially passaged until a marked reduction in proliferative capacity and an enlarged, flattened morphology consistent with cellular senescence were observed. The final passage numbers reached were passage 27 for donor 0M, passage 23 for donor 4M and passage 20 for donors 22F, 44F, and 62F.

### OSK rejuvenation

Lentiviral vectors were used to enable doxycycline-inducible expression of OCT4, SOX2, and KLF4 (OSK). Cells were transduced with lentiviruses carrying polycistronic tetracycline-controlled transcription silencer (tTS), reverse tetracycline responsive transcriptional activator M2 (rtTA) and hygromycin resistance elements (LV-CMV::tTS-T2A-rtTA-P2A-Hygro Vector builder VB010000-9369), together with the polycistronic LV-TRE::hOct4-P2A-hSOX2-T2A-hKLF4; EF1α::miRFP670nano-NLS, in which OCT4, SOX2, and KLF4 are expressed under the control of a tetracycline-responsive element (TRE).

Cells from 44F and 62F donors were dissociated and transduced at a multiplicity of infection (MOI) of 10 using spinfection at 32°C and 800 × g for 1 h, after which they were replated into flasks. Twenty-four hours post-transduction, hygromycin was added at a final concentration of 50 µg/ml to select for transduced cells, and selection was maintained for 5 days. To induce OSK expression while limiting de-differentiation, cells were subjected to two cycles of doxycycline treatment consisting of 5 days on doxycycline followed by 2 days off. Following induction, doxycycline-treated cells, no-doxycycline controls, and untransduced cells were replated into 96-well plates for downstream assays.

### Pharmacological treatments

For validation experiments using aging- and rejuvenation-associated compounds, two concentrations were tested per drug: bleomycin (35 and 75 µM), etoposide (100 and 150 µM), rotenone (1 and 2.5 µM), antimycin A (5 and 10 µM), thapsigargin (0.5 and 1 µM), MHY1485 (7.5 and 10 µM), dasatinib/quercetin (0.00005/0.1 and 0.0001/0.5 µM), trehalose (10,000 and 20,000 µM), AICAR (100 and 250 µM), and rapamycin (5 and 20 µM).

For large-scale screening, an FDA-approved library of 190 compounds (Tocris, 7200) and curated anti-aging libraries comprising 254 compounds (MCE, HY-L134; ChemFaces, L20023) were used. Main stock (10mM) plates were serially diluted using an Agilent Bravo automated liquid handling system to generate four working plates at 2× final concentrations (200, 20, 2, and 0.2 µM). Compounds were administered via half-medium exchange, with fresh medium added from the corresponding working stock plates. For each compound, the average imAgeScore across all tested concentrations was calculated and reported.

For experiments involving TRIIM trial–associated compounds, four concentrations were tested per drug: metformin (25, 50, 250, and 500 µM), dehydroepiandrosterone (DHEA; 2.5, 5, 25, and 50 µM), and recombinant human growth hormone (rhGH; 2.5, 5, 25, and 50 ng/mL). Average imAgeScore values across all concentrations were calculated for analysis.

For all pharmacological experiments, cells were exposed to compounds for 72 hours.

### DNA methylation analysis

Approximately 0.5 million cells per sample were harvested, pelleted, and snap frozen. Four biological replicates were collected for each of the 12 donors analyzed. Genomic DNA was extracted using the DNeasy Blood and Tissue Kit (QIAGEN, 69504) according to the manufacturer’s instructions, including the optional RNase digestion step. Methylation profiling was performed at the Clock Foundation (clockfoundation.org) using the Infinium MethylationEPIC BeadChip (Illumina). The “noob” background correction method was applied from the minifi R/Bioconductor package ^29^ (via preprocessNoob R function^30^) to generate b-values from the raw IDAT image data. DNA methylation age estimates were then derived using the DNAmAge general clock.

### Cell Painting and feature extraction

Cell Painting reagents were obtained from Revvity and used according to the manufacturer’s protocol. SYTO 14 (CP61) was used to label nucleoli and cytoplasmic RNA; wheat germ agglutinin (WGA; CP15551) to label the Golgi apparatus and plasma membrane; phalloidin (CP25681) to stain the F-actin cytoskeleton; concanavalin A (CP94881) to label the endoplasmic reticulum; MitoTracker Deep Red (CP3D1) to stain mitochondria; and Hoechst 33342 (CP72) to label DNA. All dyes were diluted in 5× buffer (PVDDA1), and cells were fixed with paraformaldehyde (Thermo Fisher Scientific, 28908), following Revvity’s Cell Painting protocol.

Imaging was performed on an Opera Phenix Plus high-content screening system (Revvity) operated in confocal mode, using a 20×/1.0 NA water-immersion objective with 1×1 binning and two-peak autofocus. Images were acquired using the following excitation and emission filter settings: 375 nm excitation with 435-480 nm emission, 488 nm excitation with 500-550 nm emission, 561 nm excitation with 570-630 nm emission, and 640 nm excitation with 650-760 nm emission. Six fields per well were captured, with ten z-stack planes acquired at 1 µm intervals.

Cellular feature extraction was performed using a custom segmentation and analysis pipeline implemented in Harmony software (version 5.2; Revvity). Nuclei were identified using the Hoechst channel, and cytoplasmic regions using the WGA and F-actin channels. Individual cells were segmented into five compartments: nucleus, nuclear ring, cytoplasm, cell membrane, and whole cell. Cells intersecting image borders were excluded to ensure that only fully segmented cells were analyzed. A total of 3,255 quantitative properties describing fluorescence intensity, texture, granularity, density, and spatial correlation were extracted at the single-cell level and subsequently averaged across all cells within each well.

### Data preprocessing and outlier removal

imAgeScore takes as input a modified table exported from Harmony, where each row represents an aggregated well and columns correspond to sample metadata and numeric assay-derived features. Highly constant features (near-zero variance) and features containing blank values were removed upstream in the pipeline to preserve signal and reduce input dimensionality.

To remove low-quality or anomalous wells prior to downstream analysis, we applied a correlation-based outlier filtering procedure stratified by age group. Within each age stratum, a sample-sample Pearson correlation matrix was compared across the remaining numeric features. For each sample, the mean correlation with all other samples in the same age group was calculated. Samples falling below a predefined correlation threshold (0.98) were flagged as outliers and removed. Control wells (untreated and DMSO-treated) were retained regardless of their mean correlation to ensure a consistent reference set within each age group.

### Method development

A high-level description of the model logic and evaluation protocol is provided in the Results. Full implementation details, including preprocessing, final architecture specification and parameter settings, can be found in Supplementary Methods.

### Immunocytochemistry

Cells cultured in 96-well plates were fixed with 4% paraformaldehyde (PFA), washed twice with PBS, and stored at 4 °C until staining. For immunostaining, plates were first washed with PBS and permeabilized in 0.3% Triton X-100 in PBS for 15 minutes at room temperature. Cells were then blocked in PBS containing 10% normal goat serum (NGS) and 0.1% Triton X-100 for 30 minutes.

Primary antibodies against γH2AX (Sigma, 05-636) and p62 (Invitrogen, PA5-20839) were diluted 1:300 in PBS containing 0.1% Triton X-100 and 2% NGS and applied overnight at 4 °C. The following day, plates were washed three times with PBS and incubated with appropriate secondary antibodies at 1:500 dilution (Invitrogen, A21245 and A10680), together with WGA (CP15551) and Hoechst 33342 (CP72), diluted in PBS containing 2% NGS. After three additional PBS washes, plates were imaged using an Opera Phenix Plus high-content screening system (Revvity).

### β-Galactosidase staining

Senescence-associated β-galactosidase activity in freshly fixed cells was assessed using the Senescence β-Galactosidase Staining Kit (Thermo Fisher Scientific, C10851), following the manufacturer’s instructions. The green detection marker was multiplexed with Hoechst and WGA to enable nuclear and cytoplasmic segmentation. Plates were imaged using an Opera Phenix Plus high-content screening system (Revvity).

### Live-imaging assays

Proteasome activity was measured in live cells using the BioTracker TAS2 proteasome activity probe (Sigma, SCT235) according to the manufacturer’s protocol. The probe was multiplexed with Hoechst and WGA to allow for cell segmentation, and plates were imaged using an Opera Phenix Plus system (Revvity).

Mitochondrial membrane potential and autophagic vehicle accumulation were simultaneously evaluated using a multiplexed live cell staining approach with tetramethylrhodamine methyl ester (TMRM; Thermo Fisher Scientific, I34361) and Autophagy Green dye (Abcam, ab139484), respectively. Cells were incubated at 37 °C for 30 minutes in staining buffer composed of 0.1 µM TMRM, 1× Autophagy Green dye, and 1 mg/mL Hoechst, prepared in FluoroBrite DMEM (Thermo Fisher Scientific, A1896701) and supplemented with 5% FBS (Sigma, F9665-500ML). Following incubation, cells were washed three times with 1× wash buffer supplied with the Autophagy Green kit prior to imaging. Plates were imaged using an Opera Phenix Plus high-content screening system (Revvity).

### Scratch wound assay

Scratch wound assays were performed using the IncuCyte S3 live-cell analysis system in conjunction with the 96-pin IncuCyte WoundMaker tool. Primary fibroblasts from 44F and 62F donors were seeded at a concentration of 5,000 cells/well on IncuCyte Imagelock 96-well plates (Sartorius, BA-04856) and treated the following compounds for three days: retinoic acid (1 µM), etoposide (10 µM), temuterkib (10 µM), CB049, and the combination of rhGH (25 µg/mL), DHEA (35 µM) and metformin (100 µM). Following the treatment period, uniform scratches were generated in each well using the WoundMaker tool according to the manufacturer’s instructions. Wells were washed twice with 1× PBS to remove detached cells, after which fresh media containing the corresponding drug treatments was added. Plates were then placed in the IncuCyte S3 system and imaged using the Scratch Wound scan mode at 3-hour intervals for 70 hours. Wound confluence (%) was quantified using the IncuCyte Scratch Wound Analysis Software module and used as a measure of wound closure over time.

Raw wound confluence data per timepoint were normalized such that each technical replicate was set to 1% wound confluence at time 0. Normalized values were log-transformed prior to visualization and plotted using GraphPad Prism. For statistical analysis, the area under the curve (AUC) was calculated for each treatment condition. AUC values were compared across groups using one-way ANOVA, followed by Dunnett’s multiple comparisons test to assess differences between each treatment and the untreated control.

### Image analysis

For all aging hallmark assays, including β-galactosidase and live-cell staining, image analysis was performed using Harmony software (version 5.2). Nuclei were segmented using the Hoechst channel, and cytoplasmic regions were defined using the WGA channel or, where WGA was not present, the TMRM channel. Fluorescence intensity for each dye was quantified at the single-cell level within the cytoplasmic compartment and subsequently averaged per well. For each plate, fluorescence intensity values were normalized to the mean of untreated (UNT) control wells. Statistical comparisons between compound-treated samples and DMSO vehicle controls were performed using one-way analysis of variance (ANOVA) followed by Dunnett’s multiple-comparisons test in GraphPad Prism.

## Results

### Developing imAgeScore and assessing model performance

To enable high-throughput screening of large compound libraries, we sought to establish a robust cell-based assay capable of detecting age-associated phenotypes in primary human fibroblasts with high confidence. As an initial approach, we generated fibroblast cultures from 12 donors spanning a broad range of chronological ages (0-67 years) and evaluated four canonical hallmarks of aging (genomic instability, epigenetic alterations, impaired macro autophagy, and cellular senescence) using established biomarker-based assays. Two individual readouts were assessed per hallmark, and no discernible age-related patterns were observed (Supplementary Fig. 1), suggesting that these assays may lack the sensitivity required to resolve donor age differences in vitro, or that tissue-level aging signatures may be attenuated following adaptation to culture conditions.

Given these limitations, we developed a hallmark-agnostic approach using a Cell Painting-based assay. We expanded our cohort to 24 fibroblast donors derived from individuals aged 0-87 years. Cells were plated in 96-well format, stained using the CP protocol, and imaged by high-content microscopy (Fig. 1A, B). Following image acquisition, automated single-cell segmentation and feature extraction were performed using Harmony software. Extracted features were exported in tabular format to form the input training dataset (Fig. 1A).

**Figure 1.**
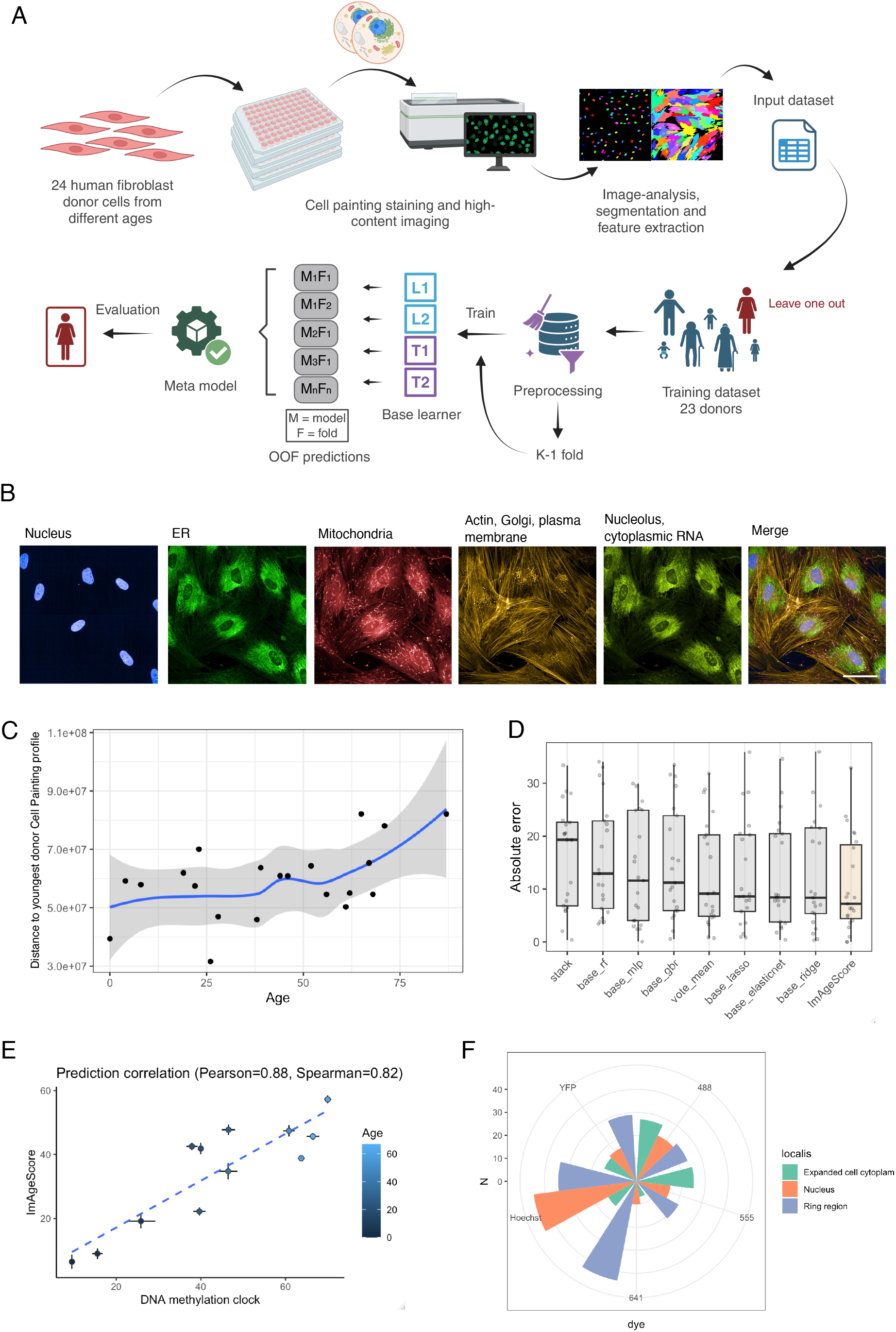
Cell Painting-derived morphological signatures predict chronological age in primary human fibroblasts. (A) Schematic overview of the workflow used to generate imAgeScore from Cell Painting data. Created in BioRender. (B) Representative Cell Painting images showing staining of major cellular compartments: Hoechst 33342 (nuclei), concanavalin A (endoplasmic reticulum; ER), SYTO14 (cytosolic RNA and nucleoli), phalloidin (actin), wheat germ agglutinin (Golgi and membrane), and MitoTracker Deep Red (mitochondria). Scale bar 50um. (C) Distance in Cell Painting feature space between each donor’s mean profile and the reference profile of the youngest donor (0M years old). (D) Absolute prediction error for left-out donors across different predictive models. (E) Correlation between imAgeScore-predicted age and DNA methylation age (DNAmAge) estimates in 12 selected donors. (F) Relative importance of the top 500 Cell Painting features contributing to imAgeScore predictions

CP profiles captured age-associated cellular variation, although the signal was distributed across many features. This was reflected by an increase in Euclidean distance between each donor’s average feature profile and that of a reference age across the cohort (Fig. 1C), indicating that chronological age is encoded in CP-derived morphological signatures. Motivated by this observation, we sought to train a predictive model capable of assigning donor age based solely on CP feature profiles.

The 24 donor samples included both technical and biological replicates distributed across multiple 96-well plates, excluding outliers (Supplementary Fig. 2A). To mitigate technical variability, features were aggregated by averaging measurements from six optical fields per well, followed by normalization to correct for plate-to-plate effects. Given the “small-n, high-p” structure of the dataset (∼1000 samples x 6000 features), we trained multiple models and evaluate performance using Mean Absolute Error (MAE) and R-squared. Model evaluation was performed using leave-one-donor-out (LODO) cross-validation to obtain per-fold predictions. To reduce overfitting, we prioritized strongly regularized models appropriate for high-dimensional data.

As no single learner was consistently optimal across donors, a stacking approach was implemented in which predictions from multiple base regressors are concatenated into a low-dimensional meta-feature matrix and used to train a regularized meta-learner. The base models were selected to provide complementary inductive biases.

We termed this final model imAgeScore. We compared imAgeScore to individual regression models, tree-based approaches, a mean voting ensemble, and an alternative stacked regressor. Feature scaling was applied to linear models, whereas tree-based models were trained on unscaled data. Across all evaluated approaches, linear models achieved the lowest MAEs (Supplementary Fig. 2D), followed by voting ensemble approaches. imAgeScore consistently outperformed all individual models, achieving an MAE of 7.23 years (Fig. 1D) and an R-squared of 0.69 (Supplementary Fig. 2C), and retained near-top performance when trained on half of the input samples (Supplementary Fig. 2B).

To benchmark performance against established aging measures, we assessed DNA methylation age using Horvath’s epigenetic clock in a subset of 12 donors. imAgeScore predictions showed a strong correlation with epigenetic age estimates (R^2^ = 0.88) (Fig. 1E), indicating concordance with DNA methylation-based clocks while capturing morphological aspects of cellular aging.

Finally, to examine the relative contributions of Cell Painting features, we quantified the importance of each stain and its associated subcellular localizations by averaging the model coefficients from imAgeScore (Fig. 1F). Features relating to mitochondrial and nuclear compartments contributed prominently to age prediction, while additional contributions arose from endoplasmic reticulum, RNA, actin, and plasma membrane associated features. These findings suggest that age-associated morphological information is distributed across multiple cellular structures rather than confined to a single hallmark.

### A method to measure biological age of artificially aged and rejuvenated cells

Replicative aging in vitro is driven by cumulative cell divisions during serial passaging and is accompanied by characteristic morphological and functional changes, including increased cell size, enhanced granularity, and reduced proliferative capacity^27,31^. To evaluate imAgeScore in this established model of cellular aging, we applied it to CP data obtained from fibroblasts undergoing serial propagation. Primary fibroblasts from five donors spanning different chronological ages were cultured until they reached replicative senescence, with the total number of passages required varying between donors. Senescence was confirmed in four donors by β-galactosidase staining (Fig. 2A). In parallel, DNA methylation analysis using Horvath’s epigenetic clock indicated advanced epigenetic ages ranging from approximately 60 to 100 years in propagated cells (Supplementary Fig. 2D).

**Figure 2.**
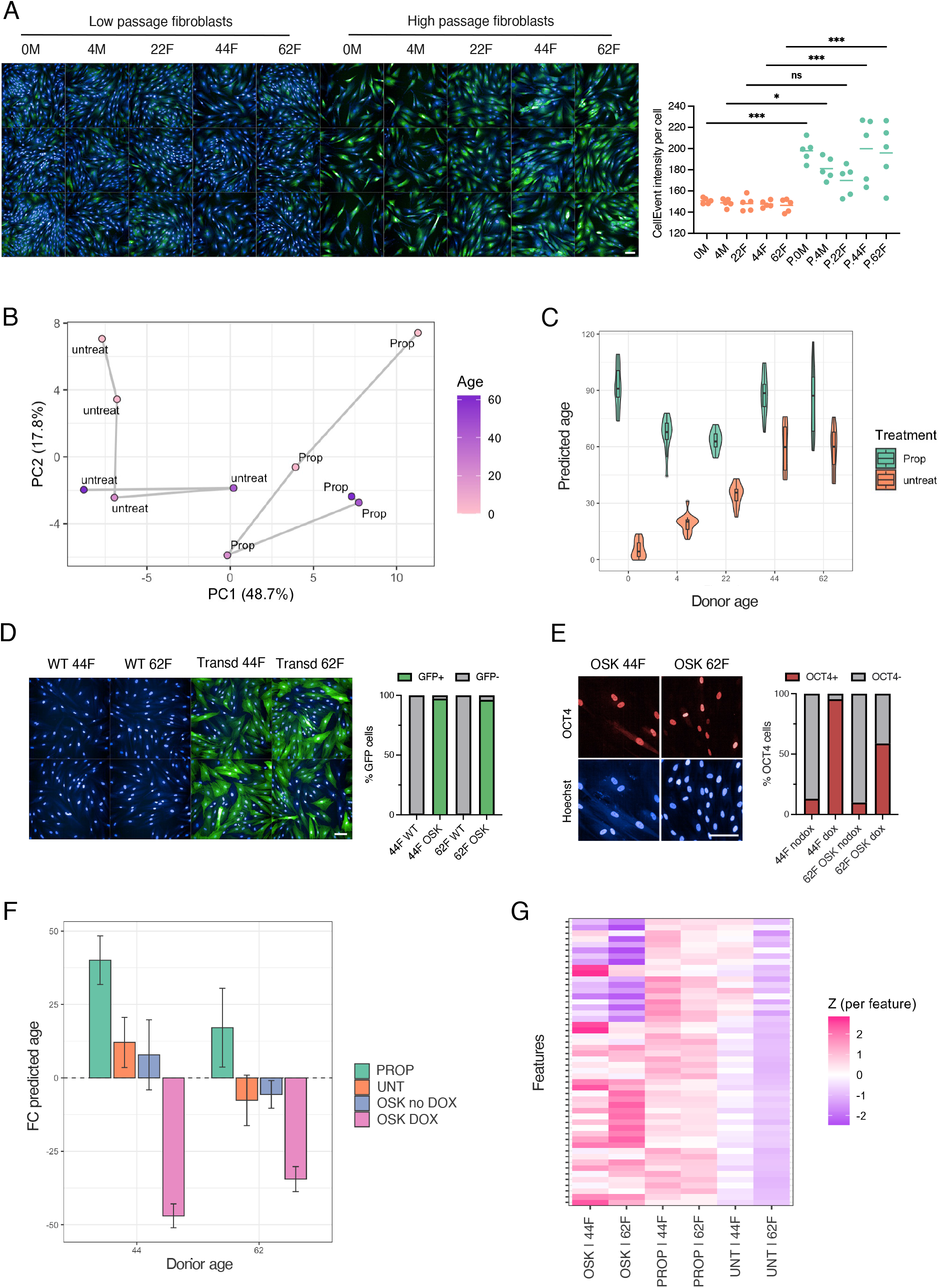
imAgeScore captures replicative ageing and OSK-mediated rejuvenation in primary human fibroblasts. (A) β-Galactosidase staining and quantification in serially propagated fibroblasts from five donors compared with low-passage controls. Scale bars, 100 μm. (mean ± s.e.m., n = 5; one-way ANOVA). (B) Two-dimensional PCA plot of the top 100 features ranked by importance, showing temporal trajectories for untreated and serially propagated fibroblasts across donors. (C) imAgeScore-predicted age in propagated fibroblasts and untreated controls for each donor. (D) GFP fluorescence–based assessment of transduction efficiency in control and transduced fibroblasts (44F and 62F). Scale bars, 100 μm. (E) OCT4 staining and quantification in OSK transduced fibroblasts with and without doxycycline treatment. Scale bars, 100 μm. (F) Fold change in imAgeScore-predicted age relative to donor chronological age (44F and 62F) in untreated, propagated, and OSK-transduced fibroblasts with and without doxycycline. Error bars represent s.e.m. (G) Z-score heatmap of the top 50 features differentiating propagated, OSK-rejuvenated, and untreated control fibroblasts, ranked by importance.

Dimensionality reduction of CP features revealed partial segregation between low-passage and propagated cells, primarily along principal component 1, while principal component 2 largely reflected baseline donor identity irrespective of propagation status (Fig. 2B). Consistent with this separation, imAgeScore detected marked age acceleration during serial propagation, with most samples ultimately predicted to correspond to approximately 90 years of age (Fig. 2C). Interestingly, donor age at the onset of propagation did not influence final imAgeScore predictions, as fibroblasts derived from 0- and 4-year-old donors reached similar predicted ages to those from 44- and 62-year-old donors (Fig. 2C), suggesting that serial propagation drives aging toward a saturation state. Notably, fibroblasts from donor 22F, which showed a non-significant increase in β-galactosidase staining relative to baseline, exhibited a more modest rise in imAgeScore, reaching approximately 60 years (Fig. 2C).

To further validate imAgeScore and assess its ability to detect cellular rejuvenation by partial reprogramming, fibroblasts from two donors (44F and 62F) were transduced with lentiviral vectors enabling doxycycline-inducible expression of OCT4, SOX2, and KLF4 (OSK; Supplementary Fig. 2G). Transduction efficiency assessed by GFP expression, approached 100% in both donors (Fig. 2D). Immunostaining confirmed robust OSK induction, with >90% of cells expressing OCT4 in donor 44F and >60% in donor 62F following doxycycline treatment (Fig. 2E).

Induction of OSK resulted in a reduction in imAgeScore-predicted age in both donors compared with untreated and no-doxycycline controls (Fig. 2F). Fibroblasts from donor 44F exhibited a near-symmetric response, with a ∼40-years increase in predicted age following propagation and a comparable magnitude of rejuvenation upon OSK induction. In contrast, donor 62F displayed a more modest age increase during propagation (∼15 years) but a larger rejuvenation response of approximately 30 years following OSK expression (Fig. 2F). The greater rejuvenation magnitude observed in donor 44F may partially reflect the higher proportion of cells successfully induced to express OSK.

Finally, we examined the top 50 features contributing to imAgeScore differences across untreated, propagated, and OSK-rejuvenated fibroblasts from donors 44F and 62F (Fig. 2G). Of these, the top 18 features were consistently elevated in propagated samples relative to controls and were reduced following OSK-induced rejuvenation, indicating that these features increase with cellular aging and are reversed upon rejuvenation (Fig. 2G). Notably, these features were enriched in measurements derived from nuclear-associated regions across multiple staining channels (Supplementary Table 2), underscoring the contribution of nuclear architecture and related processes to the aging phenotype captured by imAgeScore.

### Pharmacological validation of imAgeScore with aging and rejuvenation drugs

We next investigated whether imAgeScore could capture age-related changes induced by pharmacological perturbations using compounds previously reported to either promote or attenuate cellular aging phenotypes. Fibroblasts were treated with ten compounds targeting five distinct hallmarks of aging, comprising six damaging and four repairing interventions. Damaging compounds included antimycin A and rotenone to induce mitochondrial dysfunction^32^, bleomycin and etoposide to promote genomic instability and senescence^33–35^, MHY1485 to impair autophagy^36^, and thapsigargin to disrupt proteostasis^37^. Repairing compounds included AICAR to modulate proteostasis^38^, rapamycin to induce autophagy^39^, the dasatinib–quercetin combination to target senescence^40^, and trehalose to improve mitochondrial function^41^. Each compound was tested at four to six concentrations in fibroblasts from two donors (44F and 62F). Dimensionality reduction of CP features showed clear separation between aging- and rejuvenation-associated compounds (Fig. 3A).

**Figure 3.**
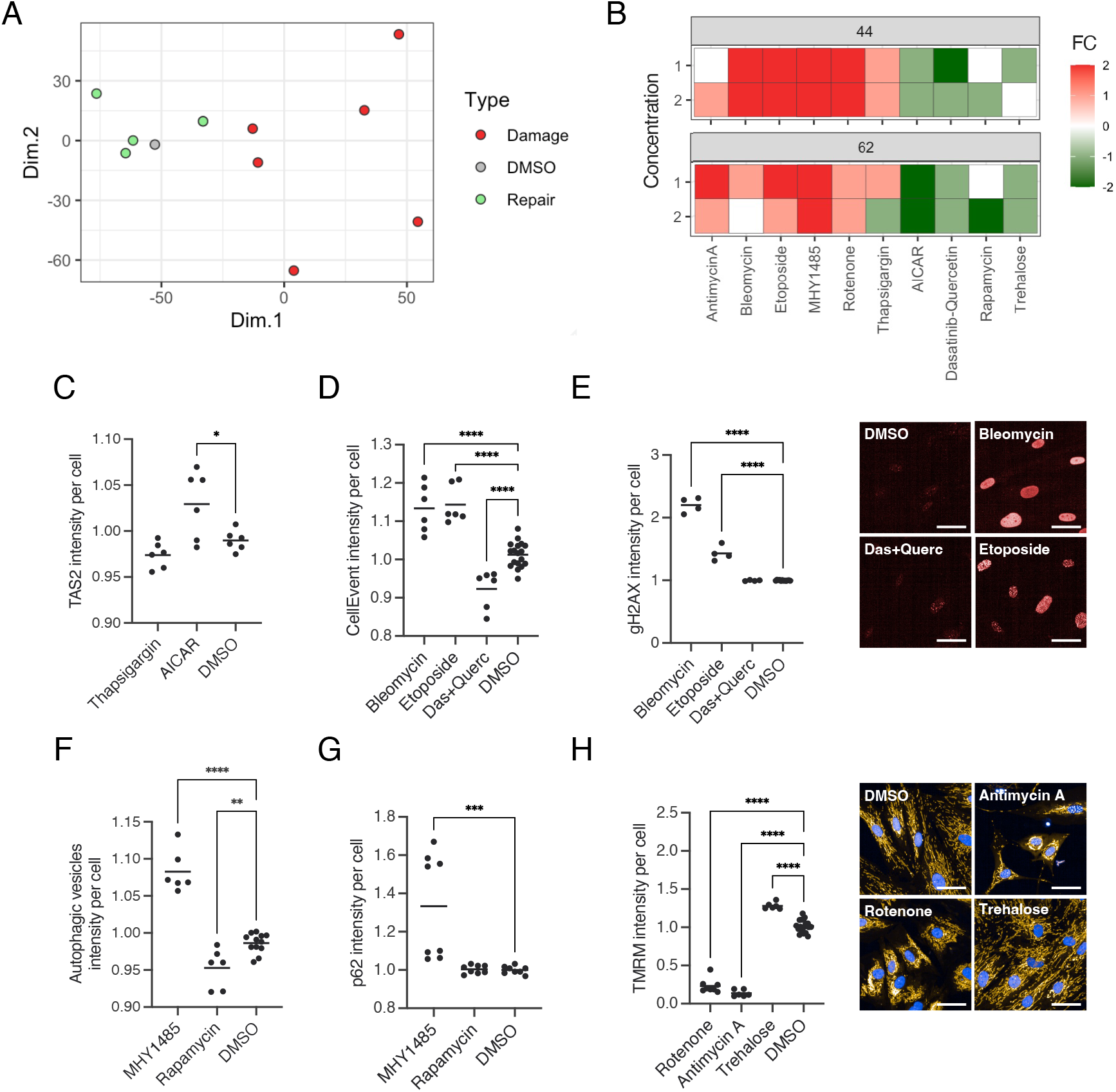
imAgeScore detects ageing and rejuvenation following hallmark-specific pharmacological perturbations. (A) Two-dimensional PCA plot of Cell Painting profiles for ageing- and rejuvenating-associated drug treatments in donor 62F. (B) Heatmap showing fold change in imAgeScore following treatment with ten ageing hallmark–modulating compounds tested at two concentrations in donors 44F and 62F. (C–H) Validation of compound-specific effects on corresponding ageing hallmarks in donor 62F using established functional assays. Proteostatic activity was assessed by TAS2 cleavage (C), senescence by β-galactosidase staining (D), DNA damage by γH2AX staining (E), autophagy by autophagosome staining (F) and p62 levels (G), and mitochondrial membrane potential by TMRM levels (H). Representative immunocytochemistry (ICC) images are shown for panels E and H. Scale bars, 50 μm. Data represent mean ± s.e.m. (n≥4; one-way ANOVA).

A logistic regression-based three-class classifier (Rejuvenation = -1, Control = 0, Aging = +1) showed strong performance when trained on a larger dataset of age-modulating damaging and repairing drugs (Fig. 3B). This approach demonstrated strong discriminability between the three classes, with AUCs of 0.978, 0.924, and 0.995 for rejuvenating, control, and damaging drugs, respectively (Supplementary Fig. 3A). The confusion matrix indicated high recall for rejuvenating and aging compounds (0.94 and 0.97, respectively), whereas controls were less separable (recall = 0.52) (Supplementary Fig. 3B). This is consistent with the expectation that rejuvenation-associated phenotypes may be subtler and partially overlap with untreated baseline states under the current feature normalization regime.

For a subset of compounds, dose-dependent effects were observed, whereby increasing concentrations of damaging agents increased imAgeScore-predicted age, while escalating doses of repairing compounds produced stepwise reductions in predicted age (Supplementary Fig. 3C). The heatmap in Fig. 3B displays two representative concentrations per compound. At these concentrations, damaging compounds consistently increased imAgeScore-predicted age, whereas repairing compounds reduced predicted age across both donors. To validate these effects, we assessed compound-specific impacts on their corresponding hallmarks of aging in donor 62F using established hallmark-specific assays. In most cases, these readouts were concordant with the direction and magnitude of imAgeScore changes observed following pharmacological treatment (Fig. 3C–H). Specifically, bleomycin and etoposide increased DNA damage and induced senescence, while dasatinib-quercetin combination reduced β-galactosidase staining (Fig. 3C,D). MHY1485 increased p62 levels, consistent with impaired autophagic degradation, and was accompanied by an accumulation of autophagic vesicles; in contrast, rapamycin maintained baseline p62 levels while reducing autophagic vesicle abundance, potentially reflecting enhanced autophagic efficiency via reduced vesicle formation (Fig. 3E,F). Rotenone and antimycin A caused a marked reduction in mitochondrial membrane potential, whereas trehalose increased it (Fig. 3G). Finally, thapsigargin showed a non-significant trend toward reduced proteostatic cleavage of TAS2, while AICAR increased proteostatic activity (Fig. 3H).

### Screening-scale application of imAgeScore identifies age-modulating compounds

To evaluate whether imAgeScore can support high-throughput discovery of age-modulating compounds, we established a screening workflow integrating automated CP and high-content image acquisition to enable reproducible and scalable profiling. Multiple concentrations per compound were tested, enabling dose-response assessment at scale (Fig. 4A).

**Figure 4.**
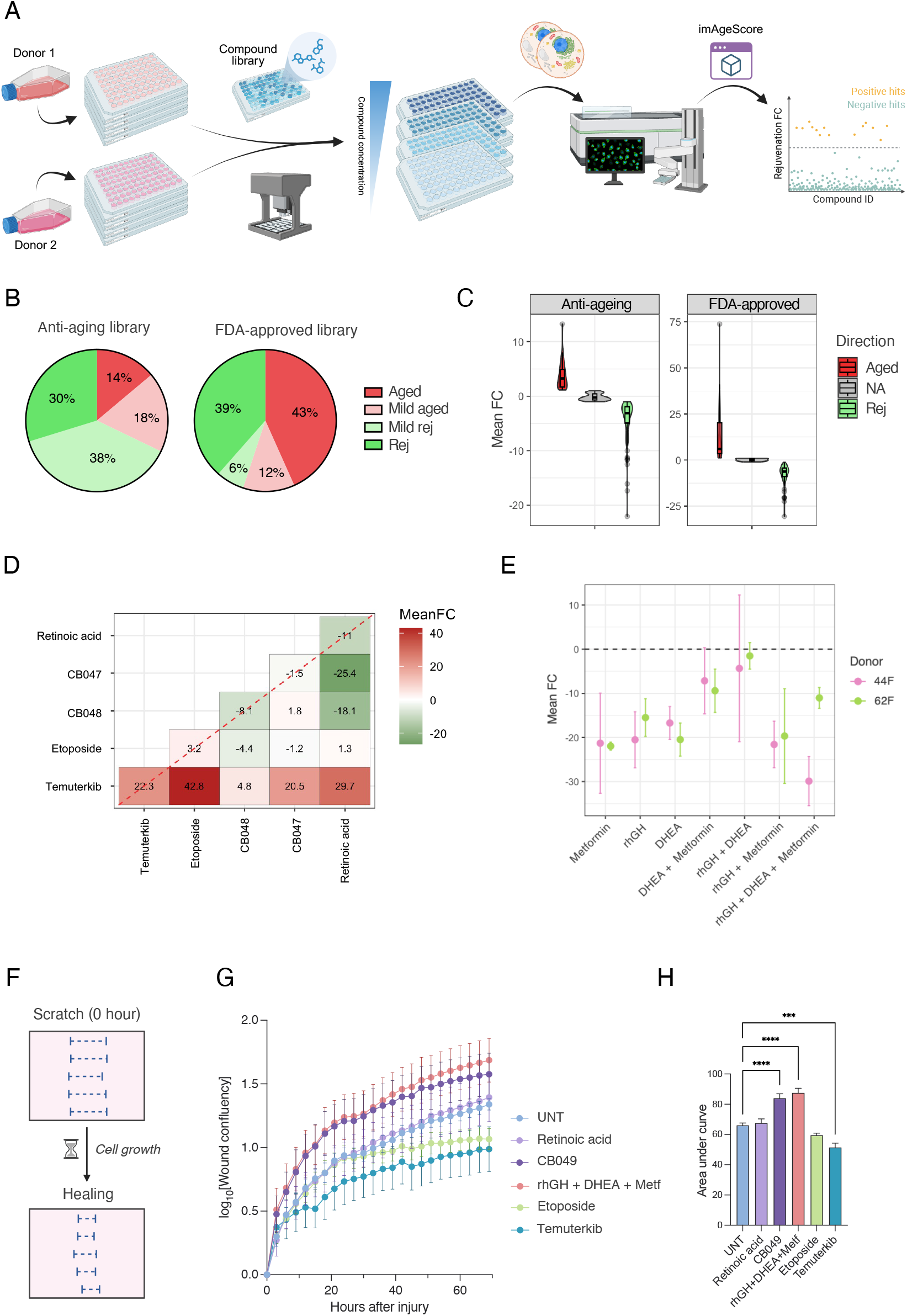
High-throughput application of imAgeScore identifies age-modulating compounds and combinatorial rejuvenation effects. (A) Schematic of the automated screening workflow integrating robotic liquid handling, Cell Painting, and high-content imaging. Multiple concentrations per compound were tested to enable dose-response profiling. Created in BioRender. (B)Distribution of fold change (FC) in imAgeScore following screening of a curated anti-ageing library of 254 compounds in primary fibroblasts from donor 62F (representative donor shown). (C) Comparative distribution of age-modulating effects across the curated anti-ageing library and the FDA-approved compound library (190 compounds), highlighting proportions of rejuvenating, ageing, and nominally neutral compounds. Compounds producing age shifts within the ±2.5-year predictive error margin of imAgeScore are classified as mild. (D) Proportion of compounds inducing concordant shifts in predicted age across the two donors following FC adjustment, evaluated across multiple FC thresholds. (E) Fold change in imAgeScore following treatment with TRIIM-associated compounds, rhGH, DHEA, and metformin, administered individually, in pairwise combinations, or as a three-compound cocktail.

We applied this approach to identify drugs with rejuvenation potential in mature primary human fibroblasts by screening two compound libraries: the Tocriscreen library of 190 FDA-approved compounds and a curated library of 250 candidate rejuvenating molecules. Screening of the curated rejuvenation library revealed that approximately one-third of compounds reduced imAgeScore-predicted age in donor 62F (Fig. 4B,C), whereas 14% increased predicted age, consistent with the intended composition of the library. Approximately half of the compounds produced only modest shifts relative to controls, with most trending toward small age reductions within the predictive margin of imAgeScore (Fig. 4B). In contrast, the FDA-approved compound library displayed a distinct distribution of effects, with roughly half of compounds increasing predicted age and 39% classified as rejuvenating. Among compounds without statistically significant effects, most trended toward age acceleration (Fig. 4B). Directionality of effects was strongly concordant between the two donors across both screens using a 2.5-year imAgeScore threshold (Supplementary Fig. 3D,E). The curated anti-aging library typically produced modest age modulation (∼3 years), with some compounds inducing changes of 10–20 years, whereas the FDA-approved library elicited more pronounced effects, averaging ∼5 years in either direction and including rare outliers approaching 75 years of predicted aging. Together, these findings indicate that a subset of candidates in the curated anti-aging library produce measurable rejuvenation in primary fibroblasts under the tested conditions. In contrast, several FDA-approved compounds reduced predicted age, highlighting potential opportunities for drug repurposing in cellular aging.

To further explore combinatorial rejuvenation strategies, we selected retinoic acid, a skin-associated rejuvenation compound identified within the FDA-approved compound library, together with two mildly rejuvenating candidates (CB047 and CB048), and assessed their pairwise combinations alongside the ageing controls temuterkib and etoposide (Fig. 4E). Combining retinoic acid with either CB047 or CB048 produced greater reductions in imAgeScore than treatment with retinoic acid, CB047, or CB048 alone, consistent with additive rejuvenation effects. In contrast, combinations including etoposide and temuterkib resulted in age acceleration comparable to single treatments; however, co-treatment with retinoic acid partially attenuated this effect, reducing the magnitude of age increase by approximately half.

We then extended our combinatorial analysis to the three compounds evaluated in the TRIIM (Thymus Regeneration, Immunorestoration, and Insulin Mitigation) trial^42^, recombinant human growth hormone (rhGH), dehydroepiandrosterone (DHEA), and metformin. We assessed these agents individually and in combination to determine whether imAgeScore detects additive or synergistic effects relative to single-drug treatment (Fig. 4F). Across both donors, most treatments resulted in reductions in imAgeScore-predicted age. In donor 44F, the three-compound combination produced the largest reduction (>20 years), consistent with an additive effect relative to single agents. In donor 62F, the three-compound cocktail induced a more modest reduction (∼10 years), comparable to or exceeded by certain individual compounds (Fig. 4F). Collectively, these findings indicate that imAgeScore can capture age-reducing effects of TRIIM-associated interventions and resolve donor-specific differences in response magnitude.

Finally, we functionally validated retinoic acid, CB049, and the rhGH/DHEA/metformin cocktail, which were selected for their pronounced reductions in imAgeScore during compound screening (Supplementary Fig. 3F). Specifically, we assessed the effects of these drugs on fibroblast wound closure following mechanical injury using a scratch wound assay (Fig. 4F). Two negative controls were also included: the senescence-inducing agent etoposide and the anti-proliferative compound temuterkib (Fig. 4G). Wound confluency increased over time in all conditions; however, area-under-the-curve analysis demonstrated significantly accelerated wound closure in cells treated with CB049 and the rhGH/DHEA/metformin cocktail. Retinoic acid did not significantly differ from control within the 70-hour observation window. As expected, both negative controls impaired wound healing, with temuterkib reaching statistical significance, while etoposide exhibited a consistent inhibitory trend (Fig. 4H). Collectively, these results functionally validate the regenerative effects of CB049 and the rhGH/DHEA/metformin combination in primary fibroblasts.

## Discussion

The ability to quantify cellular age in vitro in a scalable and biologically meaningful manner is a central requirement for the discovery of pharmacological interventions that modulate aging. Here, we describe imAgeScore, a morphology-based predictor of fibroblast age derived from high-content Cell Painting data, and demonstrate its applicability across physiological aging, replicative senescence, partial reprogramming, and pharmacological perturbation paradigms. Collectively, our findings support image-based profiling as a useful approach for measuring cellular aging dynamics and identifying candidate age-modulating compounds at scale^19,20,26,43– 46^. More broadly, these results suggest that morphology-based profiling may provide an orthogonal measure of cellular aging that complements molecular aging clocks.

While recent approaches have applied neural networks to natural image data, these methods are computationally intensive and often offer only marginal performance gains over simpler models^47^. Unlike hallmark-specific assays, which require multiple independent readouts and are not readily compatible with high-throughput workflows, imAgeScore integrates thousands of morphological features into a single quantitative metric. This hallmark-agnostic approach enabled prediction of chronological age across donors and demonstrated concordance with DNA methylation age estimates. Importantly, features contributing strongly to age prediction were enriched in nuclear and perinuclear compartments, alongside mitochondrial and cytoplasmic signals, suggesting that aging is encoded in coordinated changes in nuclear architecture and organelle organization. These findings align with accumulating evidence that alterations in nuclear morphology, chromatin organization, and mitochondrial structure are central components of the aging phenotype^48–55^. The enrichment of nuclear and perinuclear features within the imAgeScore signature further suggests that large-scale changes in nuclear organization may represent a major morphological axis of cellular aging detectable by high-content imaging.

imAgeScore robustly detected changes in cellular age across both proliferative aging and partial reprogramming paradigms. During serial propagation, fibroblasts converged toward a predicted age-saturation state, largely independent of their initial chronological age, consistent with replicative senescence representing a terminal and highly convergent morphological endpoint^56^. Conversely, inducible expression of OCT4, SOX2, and KLF4 produced substantial reductions in predicted age, with donor-specific magnitudes of rejuvenation. The reversibility of key nuclear and perinuclear features suggests that components of the morphological aging signature remain plastic and responsive to reprogramming cues. Together, these findings demonstrate that imAgeScore quantifies bidirectional transitions in cellular age state, capturing both artificial age acceleration and rejuvenation within a unified framework.

Pharmacological perturbation experiments further demonstrated that imAgeScore is responsive to interventions targeting multiple aging hallmarks. Compounds inducing mitochondrial dysfunction, genomic instability, impaired autophagy, or proteostasis disruption increased predicted age, whereas senolytic, autophagy-inducing, and mitochondrial-supporting interventions reduced it^32–41^. Concordance between imAgeScore shifts and independent functional assays indicates that the model captures biologically meaningful cellular changes rather than nonspecific morphological noise. The supervised classifier trained on damaging versus repairing perturbations further supports the separability of these states in high-dimensional morphological space.

Application of imAgeScore in high-throughput screening experiments revealed distinct patterns of age modulation across compound libraries and combinatorial interventions. Screening of a curated anti-aging library demonstrated that a subset of compounds elicited measurable rejuvenation-associated shifts in primary fibroblasts, while a minority induced age acceleration. In contrast, an FDA-approved compound library exhibited a broader distribution of effects, with substantial representation of both age-accelerating and age-reducing compounds, highlighting potential opportunities for systematic drug repurposing. Extending this approach to combinatorial treatments revealed that pairing retinoic acid with mildly rejuvenating candidates produced additive reductions in predicted age, suggesting that combinatorial pharmacological strategies may offer a promising avenue for optimizing rejuvenation therapies through additive or complementary mechanisms.

Consistent with this framework, compounds used in the TRIIM trial reduced predicted age both individually and in combination, with donor-dependent magnitudes and evidence of additive effects in certain contexts. Notably, the TRIIM intervention was originally evaluated in a clinical setting in human subjects, whereas the present study assesses these compounds within a simplified in vitro fibroblast model. The observation that the same combination produces age-reducing signatures in this cellular system suggests that imAgeScore may capture aspects of the biological pathways targeted in the clinical intervention, while also highlighting the need for further validation in more complex physiological models. Together, these findings underscore the utility of morphology-based aging metrics for stratifying and optimizing pharmacological interventions^20,46,57,58^, while also revealing inter-individual variability in rejuvenation responses^59–61^.

Importantly, functional validation using a fibroblast scratch wound assay demonstrated that selected candidates identified by imAgeScore also enhance regenerative capacity. The rhGH/DHEA/metformin combination and CB049 significantly accelerated wound closure kinetics, whereas damaging controls impaired repair. These findings support the interpretation that morphology-derived age reductions reflect biologically relevant improvements in cellular function rather than purely phenotypic shifts. More broadly, this observation suggests that high-content morphological profiling may capture functional aspects of cellular aging that extend beyond molecular biomarkers alone. At the same time, the absence of a measurable short-term effect for retinoic acid highlights the possibility that certain interventions may require longer exposure or act through mechanisms not directly captured by early wound-healing dynamics.

Several limitations warrant consideration. First, imAgeScore was developed and validated in dermal fibroblasts; its applicability to other primary cell types or more complex multicellular systems remains to be established. Second, the model exhibits a higher predictive error than established gold-standard aging clocks^62^. Accordingly, modest age-modulating effects should be interpreted cautiously unless they exceed this margin of error or are supported by independent validation. Third, pharmacological treatments in this study were administered for 72 hours, capturing short-term cellular responses. Reductions in predicted age observed over this time frame may not reflect durable rejuvenation and could, in some cases, represent transient stress responses. Conversely, certain compounds that showed limited short-term impact might exert stronger effects over longer exposure periods. Extended treatment studies will therefore be necessary to determine the stability, durability, and safety of the observed age-modulating effects. Finally, although imAgeScore responded robustly across multiple validations paradigms, its generalizability remains constrained by the relatively small training dataset. Larger and more diverse training cohorts, together with future advances in modelling approaches, may further improve predictive accuracy on unseen samples.

In summary, imAgeScore provides a practical and scalable method to quantify cellular aging and rejuvenation in vitro. Its compatibility with automated high-throughput imaging workflows positions it as a promising tool for systematic identification of compounds and combinations capable of modulating cellular age, thereby supporting efforts to develop interventions that improve human health span.

## Code and data availability

Data, code and materials will be made available upon publication.

## Acknowledgment

The authors gratefully acknowledge the Milner Therapeutics Institute and Cancer Research UK (CRUK) core facility for their support and assistance in this work.

