## Supplemental Info for "imAgeScore, a Cell Painting-Based Predictor of Cellular Age for High-throughput Drug Screening Applications"

### Supplementary Figures

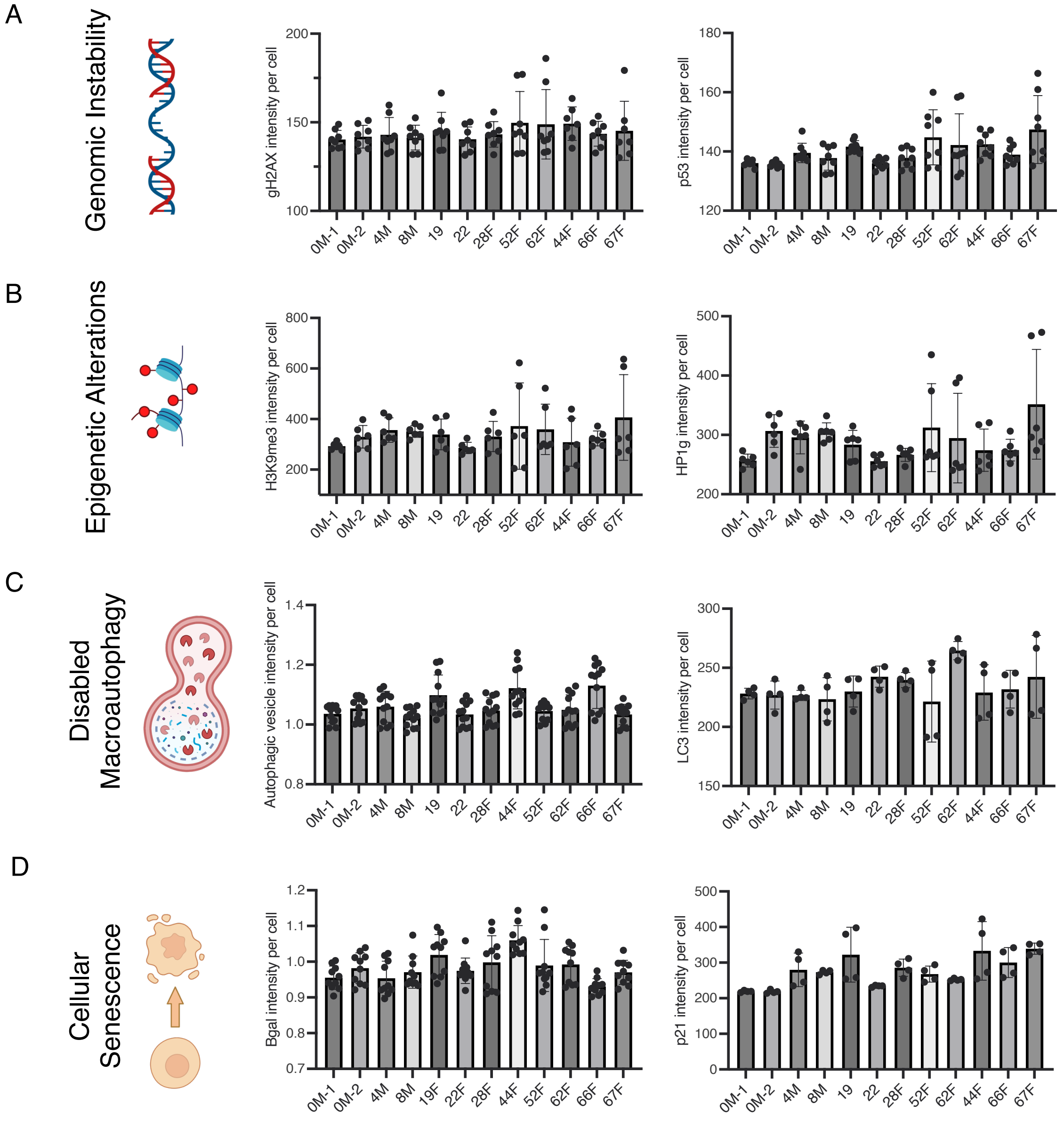


***Supplementary Figure 1. Established biomarker-based ageing assays show limited ability to resolve donor age-associated differences in vitro.***

*Assays were performed in primary fibroblasts from 12 donors spanning 0–67 years of age.*
*(A) DNA damage assessed by γH2AX and p53 staining.*
*(B) Heterochromatin evaluated by H3K9me3 and HP1γ staining*
*(C) Autophagic activity measured by autophagosome vesicle staining and LC3 immunostaining.*
*(D) Cellular senescence assessed by β-galactosidase and p21 staining.*
*Data represent n ≥ 3 replicates per donor.*
*Schematics were created in BioRender.*

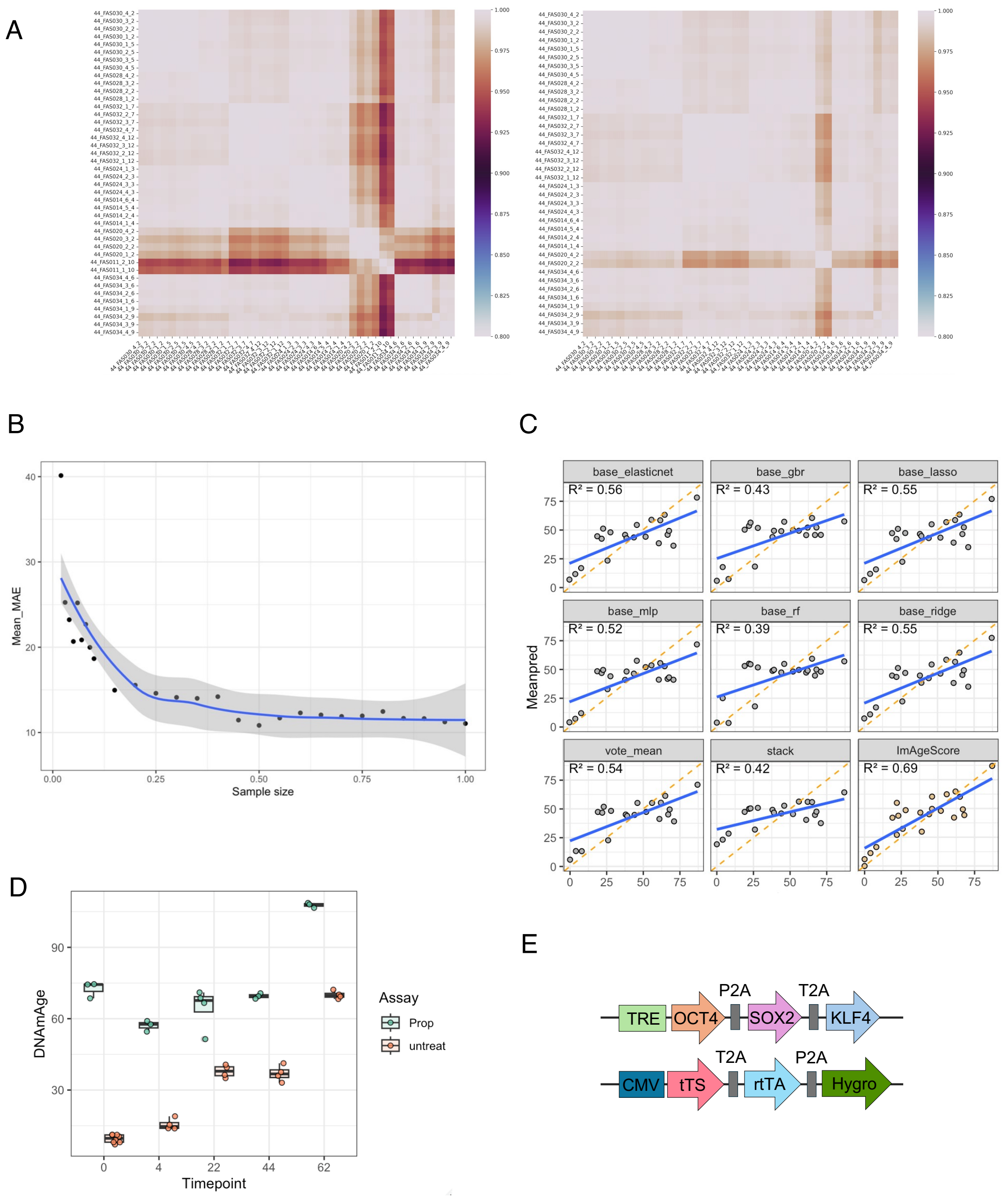


***Supplementary Figure 2. imAgeScore model development and validation in propagated and OSK-rejuvenated fibroblasts.***

*(A) Pearson correlation heatmaps between wells before (left) and after (right) outlier removal.*
*(B) imAgeScore mean absolute error (MAE) for down-sampled input datasets.*
*(C) Averaged out-of-fold predictions and $R^2$ values for different predictive models.*
*(D) DNA methylation age (DNAmAge) estimates for five propagated donor fibroblast lines.*
*(E) Lentiviral constructs enabling doxycycline-inducible expression of OCT4, SOX2, and KLF4 (OSK).*

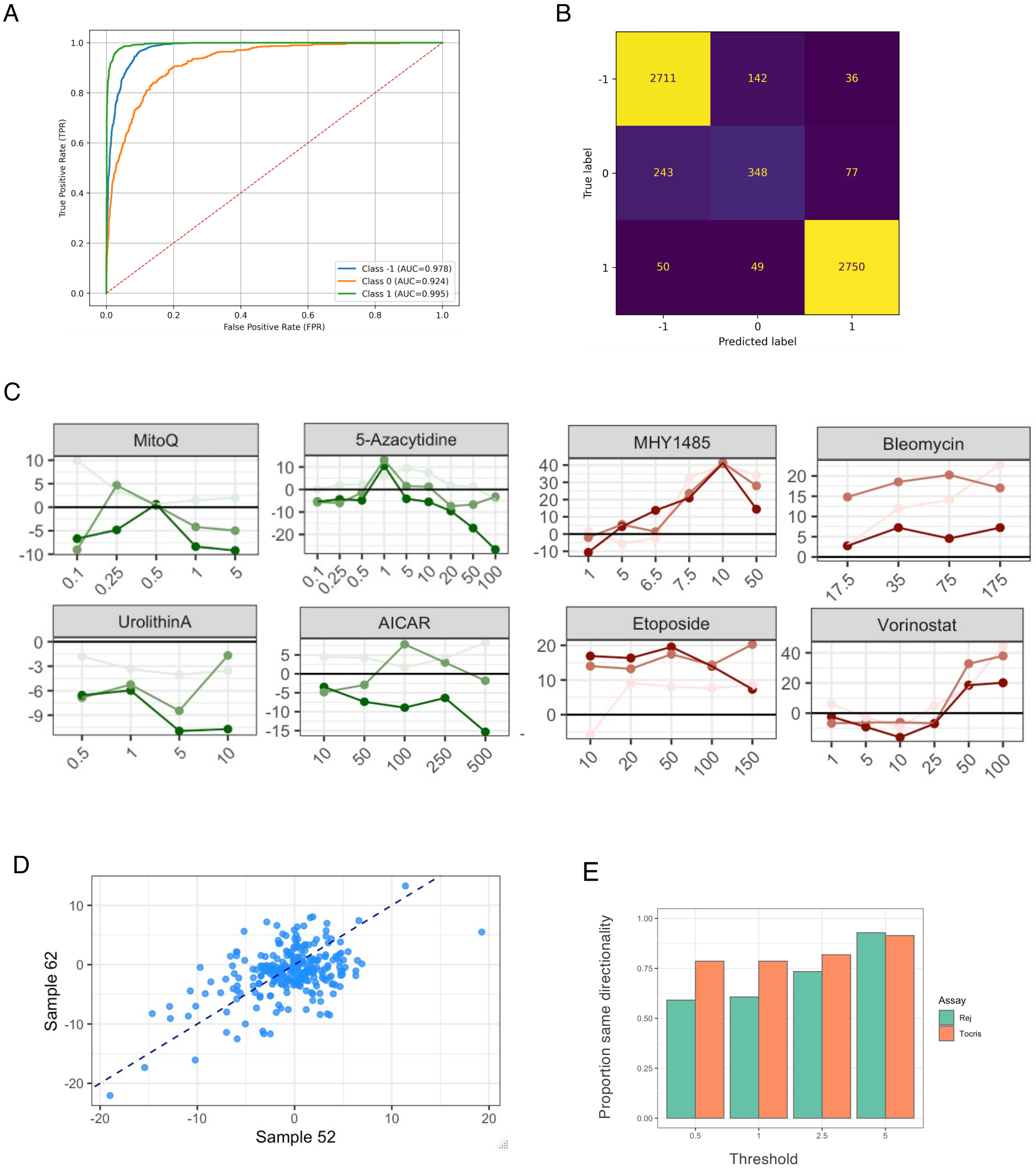


***Supplementary Figure 3. imAgeScore captures pharmacological modulation of ageing states.***

*(A) Area Under the curve (AUC) for classifier performance across three classes: -1 (rejuvenation), 0 (control), and 1 (ageing). The dashed line represents random calssification (AUC = 0.5).*

*(B) Confusion matrix showing counts of true versus predicted labels for the three classes: -1 (rejuvenation), 0 (control), and 1 (ageing).*

*(C) Dose-response curves showing predicted age changes (FC) following compound treatment relative to control conditions.*

*(D) Correlation scatterplot comparing fold change in imAgeScore for compounds from the rejuvenation screening library between donors 52F and 62F.*

*(E) Fraction of drugs that produce an age-consistent shift in the expected direction when comparing the 52F and 62F donors.*
